# Shortened lifespan and other age-related defects in bang sensitive mutants of *Drosophila melanogaster*

**DOI:** 10.1101/379321

**Authors:** Elaine R. Reynolds

**Affiliations:** Department of Biology, Lafayette College, Easton, PA 18042

**Keywords:** bang sensitive, Drosophila, mitochondria, aging

## Abstract

Mitochondrial diseases are complex disorders that exhibit their primary effects in energetically active tissues. Damage generated by mitochondria is also thought to be a key component of aging and age-related disease. An important model for mitochondrial dysfunction is the bang sensitive (bs) mutants in *Drosophila melanogaster*. Although these mutants all show a striking seizure phenotype, several bs mutants have gene products that are involved with mitochondrial function, while others affect excitability another way. All of the bs mutants (*para^bss^, eas, jus, ses B, tko* are examined here) paralyze and seize upon challenge with a sensory stimulus, most notably mechanical stimulation. These and other excitability mutants have been linked to neurodegeneration with age. In addition to these phenotypes, we have found age-related defects for several of the bs strains. The mutants *eas, ses B*, and *tko* display shortened lifespan, an increased mean recovery time from seizure with age, and decreased climbing ability over lifespan as compared to isogenic CS or *w^1118^* lines.. Other mutants show a subset of these defects. The age-related phenotypes can be rescued by feeding melatonin, an antioxidant, in all the mutants except *ses B*. The age-related defects do not appear to be correlated with the seizure phenotype. Inducing seizures on a daily basis did not exacerbate the phenotypes and treatment with antiepileptic drugs did not increase lifespan. The results suggest that the excitability phenotypes and the age-related phenotypes may be somewhat independent and that these phenotypes mutants may arise from impacts on different pathways.

## INTRODUCTION

As a cellular organelle, mitochondria are well understood, however, their role in generating system level effects that lead to malfunctioning tissues or disease is less well defined. In humans, mitochondrial DNA mutations are regularly associated with a wide range of specific defects, affecting tissues with high energy needs, such as the heart, brain, muscles, and sensorineural epithelia (Toivonen *et al*. 2001). Phenotypically, these mutations manifest themselves as hearing loss, dementia, loss of cognitive and/or motor skills, paralysis, and/or seizure. Myoclonic epilepsy associated with ragged-red muscle fibers (MERRF), have been shown to be a product of mitochondrial mutations (Wallace *et al*. 1988; Shoffner *et al*. 1990). Various neurodegenerative diseases including Parkinson’s disease have been linked to mutations in mitochondrial genes in either the mitochondrial DNA (mtDNA) or the nuclear DNA (nDNA) (Wallace, 2001). Even normal mitochondrial function damages cells over time and this accumulated damage is thought to contribute to the aging process (Askenova *et al*. 1998; Cossarizza and Salvioli, 2001; Lopez-Otın *et al*. 2016). Previous work with *Drosophila* has shown that overexpression of SOD retards oxidative damage by preventing ROS build-up that results in a longer lifespan (Sohal *et al*. 1995). Mitochondrial mutants may be even more likely to contribute to the build-up of ROS that might result in a shorter lifespan than the normal fly (Kurzik-Dumke *et al*. 1998).

Bang sensitive (bs) mutants are means to study the connections between mitochondrial dysfunction, neurological disease and aging. The bs mutants were first characterized because of their phenotype: bs mutants paralyze and seize upon receiving a short mechanical shock or “bang.” (Benzer 1971; Ganetzky and Wu 1982). Several bs mutants have been described (Table 1). When a mechanical bang is applied, these mutants first seize briefly, then remain paralyzed for anywhere between 5 and 30 seconds, then seize violently for a few seconds, and at this point are able to right themselves. Seizure is classified as coordinated contraction of all muscle sets within the fly (Pavlidis *et al*. 1995; Lee and Wu 2002). In these mutants, seizure susceptibility is enhanced as compared to wildtype (Kuebler and Tanouye 2000).

**Table 1:** Mutants that display bang-sensitive behavior and their protein product

| Gene Name | Protein Product | References |
| --- | --- | --- |
| *technical knockout (tko)* | Mitochondrial ribosomal protein | Royden *et al.* (1987) |
| *stress-sensitive B ses B* | mitochondrial adenine nucleotide translocase | Zhang *et al.* (1999) |
| *easily shocked eas* | Ethanolamine kinase mitochondrial | Pavlidis *et al.* (1994) |
| *paralyzed<sup>bss1</sup> (bss)*<br>formerly known as *bangsenseless* | An allele of the para sodium channel gene | Parker *et al.* (2011) |
| *julius caesar (jus)*<br>formerly known as *slamdance* | Unknown | Zhang *et al.* (2002a); Horne et al. (2017) |
| *knockdown kdn* | Citrate synthase | Fergestad *et al.* (2006b) |
| *bang-sensitive (bas)* | unknown | Burg and Wu (2012) |

Some of the bs genes have been shown to encode for nuclear products involved in mitochondrial function, while the other mutants do not appear, at least directly, to affect this organelle (Table 1). The bs gene *technical knockout (tko)* product is a mitochondrial ribosomal protein that affects protein synthesis in the mitochondria and *stress sensitive B (ses B)* encodes a mitochondrial adenine nucleotide translocase (Royden *et al*. 1987; Shah *et al*. 1997; Zhang et al. 1999). The *knockdown (kdn)* gene encodes citrate synthase (Fergestad *et al*. 2006b). Another bs gene, *easily shocked* (*eas*) gene encodes ethanolamine kinase, an enzyme involved in de novo synthesis of phosphotidyl-ethanolamine, which occurs within the mitochondria (Pavlidis *et al*. 1994). Decreased ATP levels have been observed in *eas, ses B* and *tko* mutants (Fergestad *et al*. 2006b). Other bs mutant do not appear to affect mitochondrial function directly. The bs mutant *paralyzed^bss^* (abbreviated as *bss* for the remainder of this paper) is an allele of the Na channel gene, while the cellular role of *julius cesear* remains unclear (Parker *et al*. 2011; Horne *et al*. 2017). However, all of these strains of bs mutants show very similar, if not identical, neurological phenotypes. The behavioral data leads to the idea that all the bs mutations could affect mitochondrial function, structure, or location within the organism. Several investigators have asked if other mitochondrial enzymes produce bs phenotypes, but genes for glycerol-3-phosphate dehydrogenase, arginase, and proline oxidase and other Kreb cycle enzymes do not have alleles that show bs phenotypes (Wang *et al*. 2004; Fergestad *et al*. 2006b).

In this paper, we document age-related aspects of a subset of the bs mutants and explore the relationship of aging phenotypes to their seizure behavior. We looked at the lifespan of bs flies, evaluated their seizure phenotype with age, and looked at their climbing ability. Defects in climbing ability have been correlated with neurodegeneration (Feany and Bender 2000). We also attempted to rescue the age-related phenotypes with melatonin and anti-epileptic drugs (AEDs). Our data suggests that the neurological phenotype and age-related phenotypes are somewhat separable. The antioxidant melatonin relieves age-related declines in some mutants, while having no impact on epileptic seizure in young flies. We also observed that AEDs decreased seizure frequency, but have no effect on lifespan. These experiments lead to a better understanding of tissue-specific dysfunction and disease.

## MATERIALS AND METHODS

### Fly Stocks

All stocks were maintained on a standard yeast, cornmeal, molasses food at 21-25°C unless otherwise noted. Flies were transferred at regular intervals to maintain continuous cultures of the strains used. For all experiments, flies were kept in a humidified, temperature-controlled environment at 25°C throughout and kept on a 12 hr light/dark schedule (Rogina, Benzer, & Helfand, 1997). Stocks for *tko, bss f* and *eas f* were originally isolated on a CS background and have been isogenized with CS through 10 generations. CS was used as the control for these strains. We also generated *w^1118^ tko, w^1118^ bss f* and *w^1118^ eas f* stocks and these stocks were used to confirm some behavioral data and for the cytochrome oxidase assays described below. Stocks for *jus* and *ses B* originated from a *white* background and were compared to *w^1118^* stock as a control. For all of these genes, there is only one allele that displays the seizures and age-related defects, although they do so in different genetic backgrounds. All stocks are negative for Wolbachia infection as determined by PCR analysis (Grandison *et al*. 2009). All stocks and other reagents used in these experiments are included in a Reagent Table as a supplemental file (see Reagent and Data Availability below).

### Lifespan Study

All experiments looking at the lifespan of a population of flies were conducted in the same general fashion. Ten female and ten male flies were transferred to plastic bottles containing a standard food, with several grains of yeast added on top and allowed to lay eggs for 5 days. From these cultures, 100 adult flies were collected from 0-5 days and maintained in a bottle containing standard food. Survivors were transferred to new food within every 5-day interval and dead flies were counted. There were at least three replicate bottles for each experiment. Mean survival percent and standard deviations were calculated for each strain from data points within the five-day age brackets. Pairwise comparisons were made with controls using T tests with significance at p < 0.05.

### Phenotype over lifespan study

Ten female and ten male flies of the strain to be tested were transferred to standard food with several grains of yeast added and allowed to lay eggs for 5 days. Young hatching flies were collected using anesthesia at 5 days of age and maintained in groups of 15-20 flies in a vial containing standard food. During every 5-day interval of the lifespan, the flies were placed into a clean empty vial and rested for 20 min. Bang sensitivity was measured by vortexing each vial of flies for 10 sec at a maximum speed. Recovery time for individual flies was then recorded. Data was collected from at least 3 replicate vials in each experiment and the whole experiment was repeated twice. Mean recovery time and standard deviations were calculated for each strain from data points within the five-day age brackets. Comparisons were made with controls using T tests with significance at p < 0.05.

### Climbing Assays

Flies used in the climbing assay were raised and maintained as described for the phenotype study. During every 5-day interval of the lifespan, the flies were placed into a clean empty vial and rested for 20 min. Flies from each vial then gently tapped to the bottom of the vial. The number of flies that climbed above a line 4 cm from the top of the vial was counted after 20 sec (Feany & Bender 2000). Occasional, especially later in the lifespan, a few flies would paralyze due to the tap. However, the number that paralyzed in any given experiment was small and these flies were excluded from that climbing data point. The process was repeated 5 times with a two-minute rest period between trials to determine an average for the data point. For initial experiments, after allowing the vials to rest for 20 min, the climbing assay was repeated. The second assay was dropped in later experiments after it was determined that the data collected at both times were statistically the same. The mean percentage of flies to cross the line in 20 seconds and standard deviations for each time point were calculated for each strain. Strains at various time points were compared using T tests with significance at P < 0.05.

### Melatonin and antiepileptic drugs studies

Ten males and ten females were placed into a plastic bottle standard food containing 100μg/ml melatonin (Sigma, St. Louis, MO) and allowed to lay eggs for 5 days. A control culture was also raised without the drug. At 5 days of age, flies reared on the melatonin-infused food and the control were collected and placed in plastic bottles containing the same food mixture. The effect of melatonin as compared to the control on lifespan, phenotype and climbing ability of the bs strains was determined using the procedures outline above.

In a similar fashion the effect of the antiepileptic drugs phenytoin and valproate was tested on the lifespan of one bs strain, *easily shocked (eas)*. Various doses of the drugs were examined by raising both CS and *eas* in standard fly food infused with the drugs. Flies were maintained on the drug during their lifespan. These doses had previously been shown to reduce the antiepileptic phenotype of this strain at 5 days of age as evidence by a reduction in the MRT and the percentage of flies paralyzed (Kuebler and Tanouye 2002; Reynolds *et al*. 2004). For each drug and each dose, three replicate bottles were performed and the experiment was repeated three times. Lifespan data was collected and analyzed as described above.

### Seizure and lifespan experiments

Experiments looking for an effect of seizure on lifespan were conducted in the same fashion as other lifespan experiments except that the initial number of flies per bottles was 40. Each day during the lifespan study, flies were transferred without anesthesia to a clear vial and then vortexed for 10 seconds. After recovery, flies were then transferred back to the holding bottle. Lifespan data was collected and analyzed as described above.

### Mitochondrial Isolation and Cytochrome C Oxidase Assay

Flies were collected either 5 days (young) or 15 days (old) after eclosion. One hundred flies were homogenized in 1.5 mL of PBS using a Tissue Tearor. Mitochondria were isolated per kit instructions (Thermoscientific, Rockford, IL). The pellet was resuspended in 200 ul of PBS then sonicated on ice. The samples were stored at −80° C until testing. Cytochrome C oxidase is an enzyme associated with the electron transport chain of mitochondria. Assays were preformed as directed by a cytochrome C oxidase assay kit (Sigma-Aldrich, St. Louis, Missouri) as per instructions. Briefly, reduced cytochrome C was generated by reducing Ferrocytochrome C (FCC) by the addition of DTT and incubation until the 550/565 ratio was at least 10. Each assay contained appropriate amounts of assay buffer, enzyme buffer, reduced cytochrome C, and 50 ul of the mitochondrial isolate. All absorbance was read immediately at 550nm for the first 45 seconds of the reaction. Enzyme activity was calculated by looking at the change in absorbance over the time interval and using the change in extinction coefficient between the oxidized and reduced state to estimate the amount of activity per ul of sample. Activity was reported as U/mg based on a standard BCA protein assay (Markwell *et al*. 1982). At least three assays were preformed independently for each strain and time point. Comparisons were made with controls using T tests with significance at p < 0.05.

### Reagent and Data availability statement

All reagents and data from this work are available. The reagents used including fly stocks and kits are listed in the Reagent Table provided as a supplemental file FileS1. Fly stocks are available from the author. All the data collected for these experiments is available as supplemental file FileS2. Supplemental files available at FigShare.

## RESULTS

### Lifespan Study

The mitochondrial mutants, *tko* and *ses B*, both showed a dramatic reduction in lifespan versus the control strains (Figure 1A, Cs is shown, CS-*w^118^* p >0.1). By 30 days of age, every fly in the *ses B* population had died, while the wildtype had 70±15.2 % of its population remaining. The *tko* flies had no survivors by 45 days of age, while CS had 42±6 % of the population alive. Both the *tko* and *ses B* flies showed a significant reduction in the survival percent by 11-15 day range (CS-*tko* p=0.0027, CS-*ses B* p=0.0011). The *eas* strain also showed a decrease in lifespan as compared to the control with significance being reached at 21 days of age (p=0.0426) and with no survival past 55 days of age (Figure 1A). The *bss* or the *jus* population, however, did not show a significant reduction in lifespan versus the control at any age. Comparisons were also made with *tko, eas* and *bss* mutants crossed into a *w^1118^* background. The same results were observed; *tko* and *eas* showed reduced lifespans compared to the *w* control (data not shown).

**Figure 1.**
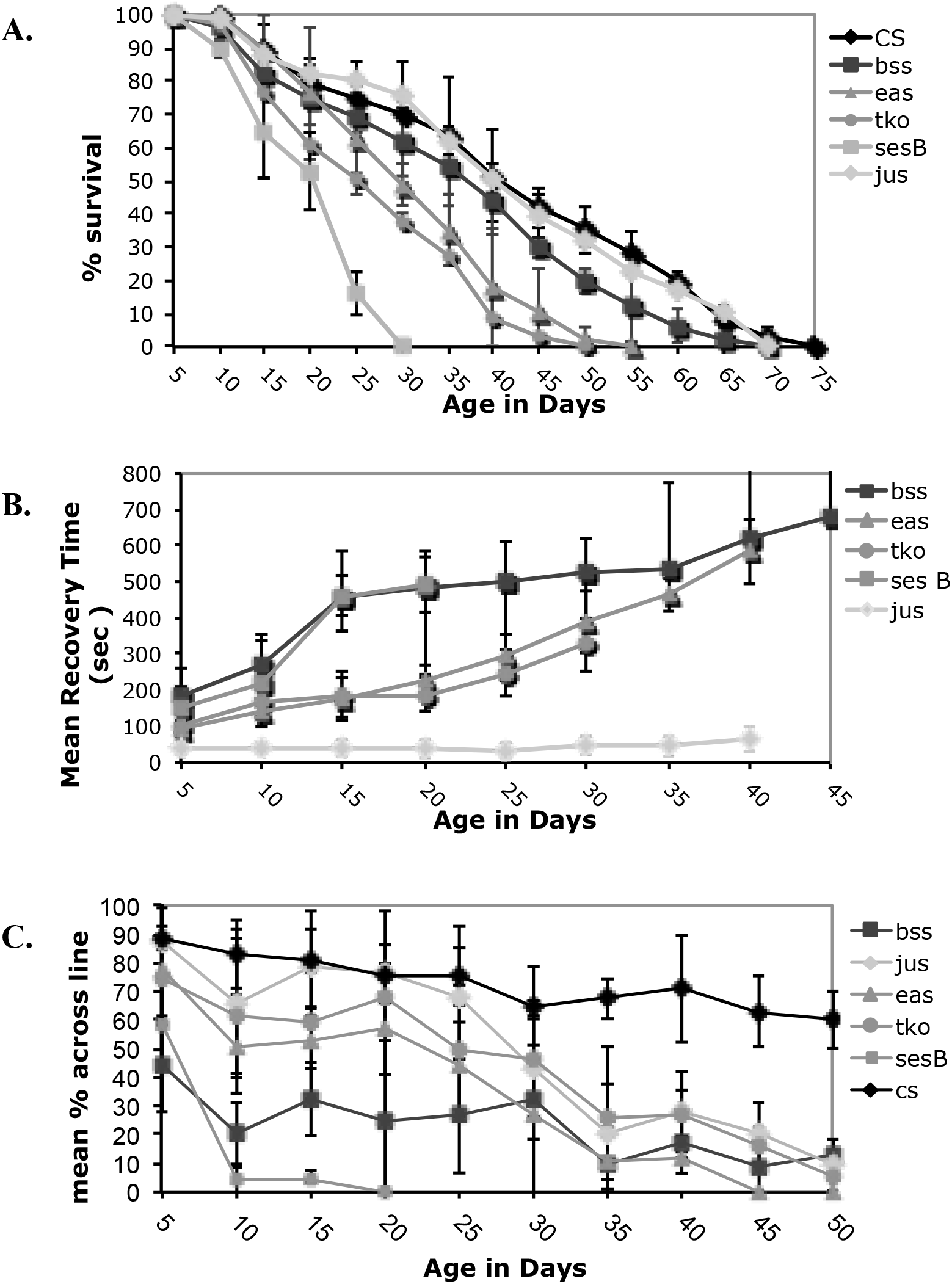
Lifespan and age-related phenotypes for bang sensitive mutants. A) Lifespan B) Mean Recovery Time C) Climbing ability. All error bars are presented as standard deviations.

### Other age-related phenotypes

Over their lifespan, all of the bs strains except *jus* showed a dramatic worsening of the bang-sensitive phenotype, as characterized by an increase in mean recovery time (Figure 1B). The decline occurred at a similar rate for the affected strains. The *eas* strain showed the largest change with age, beginning with a 94±25 s recovery time and ending with a 585±90 s recovery time at 41 days of age (p <0.001). The *ses B, tko* and *bss* strains also showed increases in recovery times: *ses B* from an initial 151±56 s recovery time to 491±74 s recovery time by 16 days old; *tko* from an initial 100±30 s recovery time to 327±75 s recovery time by 35 days of age; and *bss* from a 188±65s initial recovery time to a 658±209 s recovery time at 36 days of age. The control CS or w strains do not exhibit a bang-sensitive phenotype over their lifespan.

All mutants also showed a significant decrease in climbing ability with age (Figure 1C). The mutant strains *ses B* and *bss* were found to have a decreased ability to climb from an early age. All other mutants demonstrated initial climbing abilities equivalent to the wild type. By 30 days of age, all bs flies showed a significant reduction in climbing ability as compared to the normal fly (p<0.001). Wild type flies showed a gradual decrease in climbing ability with age.

We also look at cytochrome oxidase activity over age for *w*, and *bss, tko* and *eas* mutant stains in a *w* background (Table 2). We found that *tko* and *bss* had significantly higher levels of cytochrome oxidase in both 5 and 15 day old flies (*w* vs. *tko* P<0.004 for both, *w* vs. *bss* P<0.02 for both), but the mutant *eas* had no significant difference in cytochrome oxidase activity. Over that time frame, there was no difference in cytochrome oxidase with age for the control or mutant strains.

**Table 2.**
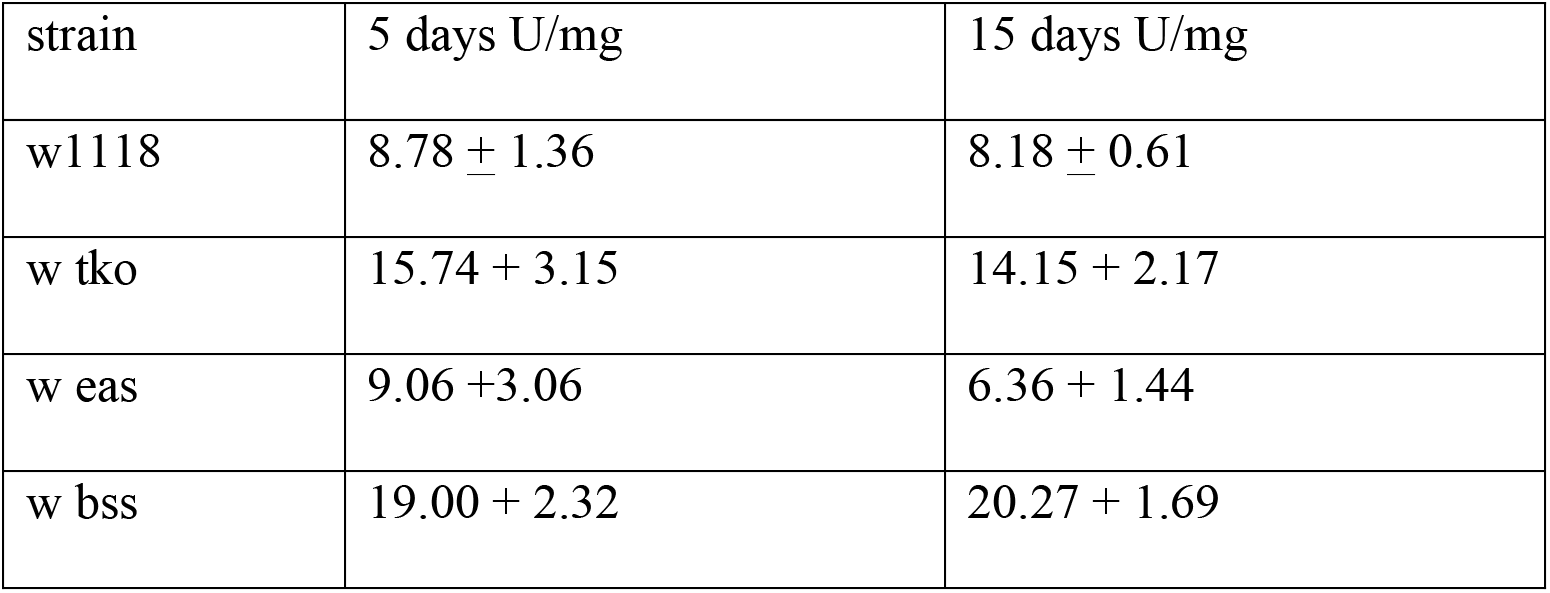
Cytochrome Oxidase Data at 5 and 15 days

### Effects of melatonin on age-related phenotypes

In separate experiments, bang sensitive mutants that showed a reduction in lifespan (*tko, ses B, and eas*) were raised with melatonin, a known free radical scavenger, in an attempt to investigate whether the age-related phenotypes could linked with increased ROS in the mutants. The melatonin increased the lifespan in all mutant stains, except *ses B* (Figure 2A). For example, *tko* without melatonin flies were all dead by the age of 45, while 45±1.2 % of the *tko* melatonin flies were alive at that same age. The *eas* mutants raised with melatonin had a 55±2% survival at 50 days of age, while all *eas* flies raised with out melatonin were dead by 50 days of age. The melatonin did not increase the lifespan of the *ses B* mutant as compared to the *ses B* flies that were raised without the melatonin (p>0.8 for all time points). CS flies grown on melatonin did not show a significant difference in lifespan.

**Figure 2.**
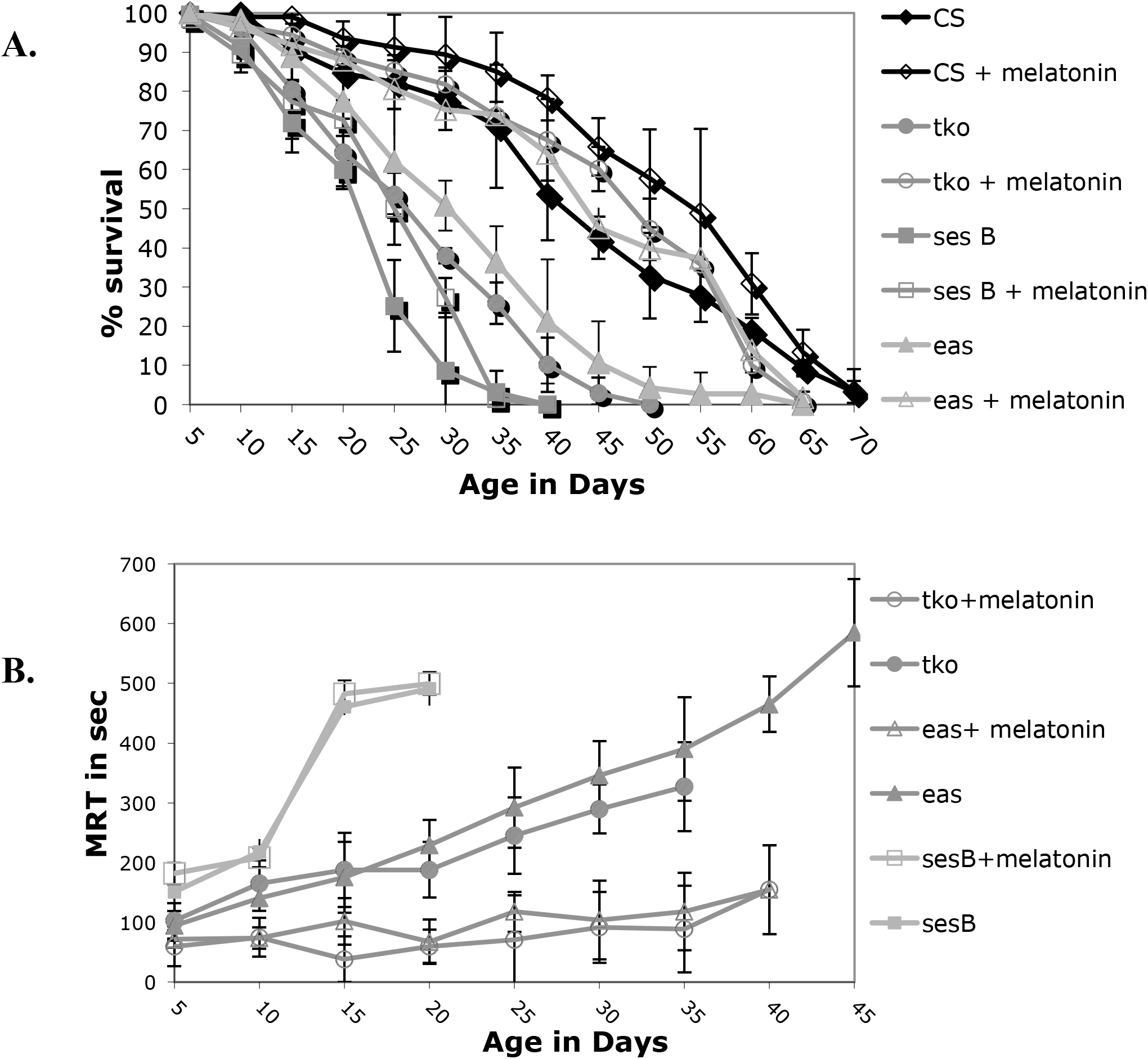
Lifespan and age-related phenotypes for selected bang sensitive mutants with melatonin supplementation. A) Lifespan B) Mean Recovery Time. All error bars are presented as standard deviations.

Melatonin supplementation significantly reduced the effect of age on recovery time for the *tko* and eas mutants but not for *ses B* (Figure 2B). The initial recovery time at 5 days of age for control *tko* mutants was 100±30 s, while the initial recovery time for melatonin raised *tko* flies was 60±31 s. By the 35 day age bracket, the control *tko* flies took 327±41 s to fully recover, while the melatonin *tko* mutants only took 88±72.5 s to recover during the same age range. The melatonin raised *eas* population also showed a striking reduction in recovery time, especially later in their lifespan (p=0.0114). By 40 days of age, the control *eas* flies took an average of 465±46 s to recover, but the melatonin *eas* population needed an average of 154±65 s for recovery. The melatonin-supplemented *ses B* and control *ses B* strains did not show any significant difference in recovery times at any point in their lifespan (p=0.1335). Melatonin supplementation also had an impact on recovery time on the *bss* mutant, which previously was shown to have an increasing MRT with age (data not shown in Fig 2). At 40 days of age, the control *bss* population had a mean recovery time of 658±209 s; in contrast, the melatonin-raised population had a mean recovery time of 145±44 s at 40 days old. While melatonin supplementation did improve significantly the initial MRT of several mutants, seizure was not completely eradicated in any bang sensitive mutant. These melatonin results suggest that some bs mutants might have age-related phenotypes mediated by ROS, but that some mutants like *ses B* cannot simply be rescued by a reduction in ROS levels.

### Effects of Anti-epileptic drugs on age-related phenotypes

It could be that the hyperexcitability produced by bs mutants leads to degeneration or other declines in nervous system function that result in the reduced lifespan. We designed two different experiments to look at the relationship between hyperexcitability or seizure and decreased lifespan. First we asked if generating a seizure everyday using mechanical shock could lead to a decreased lifespan. While there was a slight decrease in lifespan with daily mechanical stress, the decrease was also observed in wildtype and is likely to physical damage accumulated by vortexing (Figure 3A).

**Figure 3.**
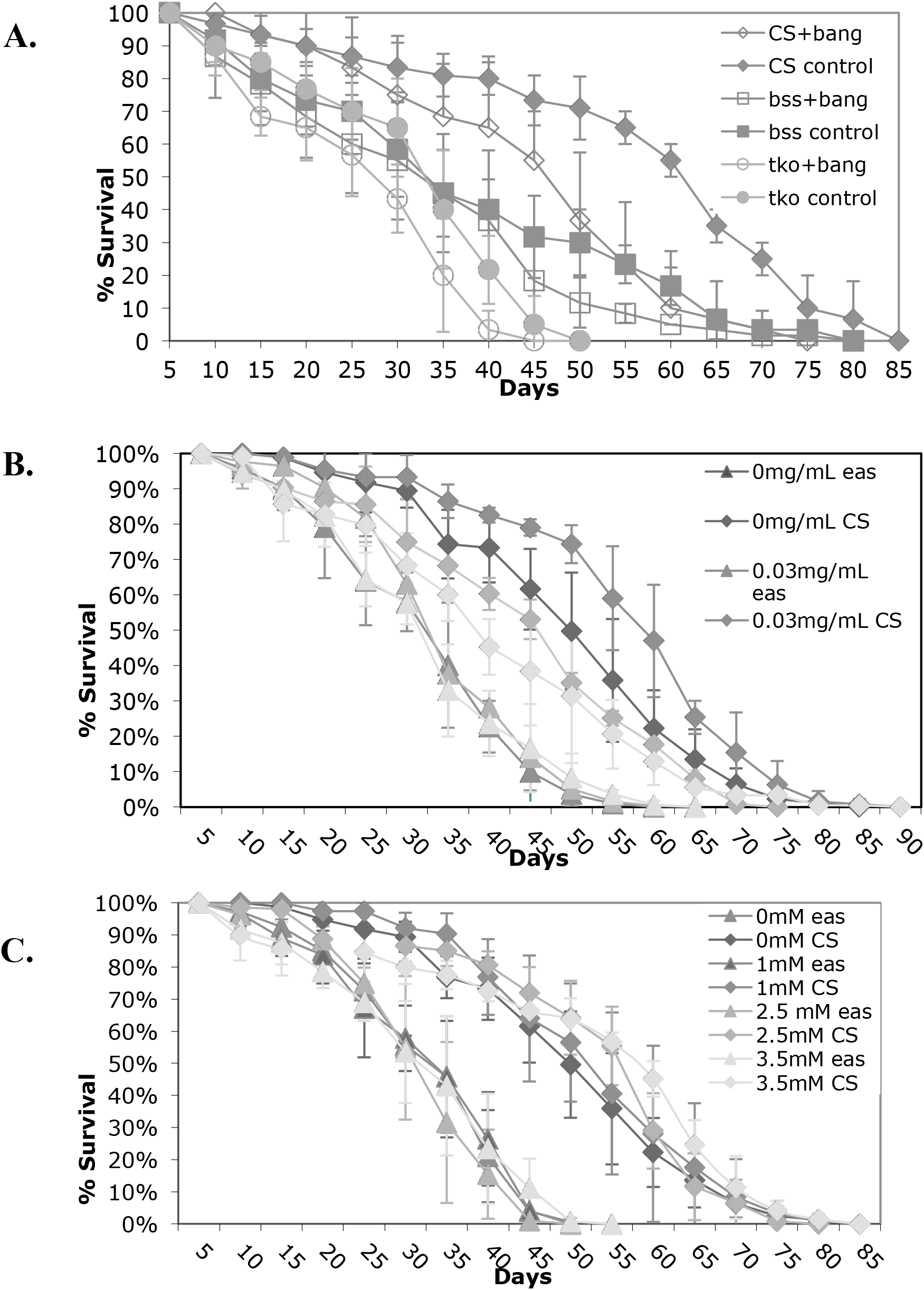
Decreased lifespan is not related to epileptic phenotype. A) Flies were vortexed every five days over lifespan B) Flies were treated with phenytoin as indicated. C) Flies were treated with valproate as indicated. All error bars are presented as standard deviations.

In previous work, it had been shown that several antiepileptic drugs decreased the percentage of seizure in bs mutant flies as well as the time necessary to recover from seizure (Kuebler and Tanouye 2002; Reynolds *et al*. 2004). We treated the bs mutants with various concentrations of the antiepileptic drugs phenytoin and valproate for their effects on lifespan. Surprizingly, the drugs had very little effect on lifespan. Figure 3B and C show the lifespan data for phenytoin and valproate respectively for the *eas* mutant, which had shown the strongest drug effect in previous work. The administration of phenytoin and valproate throughout development at several doses did not rescue the defects in lifespan for *eas* mutants. Overall, these results do not support the hypothesis that the behavioral phenotype and the age-related defects are functionally related.

## DISCUSSION

This study found that lifespan was altered in several bang-sensitive mutants, and that the seizure phenotype and climbing ability became worse with age. A summary of the age-related phenotypes of the mutants is presented in Table 2. Previous studies have demonstrated that both the *tko* and *ses B* strains have mutations in the gene that encodes for nuclear mitochondrial products (Royden *et al*. 1987; Shah *et al*. 1997; Zhang *et al*. 1999). The vital role that mitochondria play in the basic energy production in all tissues posits that any mitochondrial defect might produce physiological and behavioral changes. The defects in mitochondrial mutants *tko* and *ses B*, in addition to bs behavior, produced a reduced lifespan, worsening of the bang-sensitive phenotype with age, and a decrease in climbing ability with age. The *tko* mutation affects a mitochondrial ribosome protein and has a broader impact on mitochondrial function, while the *ses B* mutation affects the mitochondrial adenine nucleotide translocase genes. Some authors have suggested that both mutants have strong impacts on energy production (Fergestad *et al*. 2006b; Kemppainen *et al*. 2016). Toivonen *et al*. (1999) show that *tko* impacts a number of mitochondrial complex enzymes in contrast to our study, which finds an slight increase in cytochrome oxidase function as compared to control. Our findings are supported by another study which show that mitochondrial mutants have been shown to compensate for their energetic losses (Celotto *et al*. 2011). In addition, Kemppainen *et al*. (2014) found that *tko* is not rescued by an alternative oxidase, which should alleviate cytochrome oxidase insufficiency. This suggests that *tko* is a complex mutant with multiple metabolic effects. Our experience is that the phenotypes (both seizure and age-related) and perhaps mitochondrial function in these mutants is strongly tied to diet and or background and these differences may have created the observed differences. Another interesting finding of this study was that melatonin was able to rescue the *tko* age-related lifespan and phenotypic effects, but not the *ses B* age-related defects. Melatonin does not rescue the bang-sensitivity of either mutant. This also supports the idea that *tko* has broader effects than *ses B*, which may lead to a number of defects related to both energy and ROS production, but that age-related impacts on lifespan and other phenotypes are not simply due to increased ROS production in mutants.

**Table 2.** Summary of characteristics for each bs strain

|  | **CS/w** | ***tko*** | ***ses B*** | ***eas*** | ***jus*** | ***bss*** |
| --- | --- | --- | --- | --- | --- | --- |
| Seizure phenotype |  | STRONG | WEAK | STRONG | WEAK | STRONG |
| Shortened Lifespan | NO | YES | YES | YES | NO | NO |
| Decreased Phenotype | NO | YES | YES | YES | NO | YES |
| Decrease in Climbing Ability | NO | YES | YES | YES | YES | YES |
| Melatonin Lengthens Lifespan | NO | YES | NO | YES |  |  |
| Melatonin Rescue Phenotype | NO | YES | NO | YES |  | YES |
| AED rescue |  |  |  | NO |  |  |

Blank indicates Not Done

The *eas* mutant also showed a decrease in lifespan, worsening of the bs phenotype, and a decrease in climbing ability. From this data, it seems likely that the *eas* mutant may impact mitochondrial function like *tko* and *ses B*. Previous work shows that *eas* encodes an ethanolamine kinase involved in membrane phospholipid synthesis located in the mitochondria (Pavlidis *et al*. 1994). Mutants have been shown to have abnormal phospholipid composition, especially in the fly head (Nyako *et al*. 2001). This mutant has also been shown to have reduced ATP levels although not as decreased as *tko* and *ses B* (Fergestad *et al*. 2006). Melatonin also rescues the *eas* phenotype, indicating that the phenotype might be complex as well, as might be expected for a mutant that alters phospholipid composition. Another mutant, *enigma*, which encodes a mitochondrial enzyme involved beta-oxidation of fatty acids and lipid homeostasis extends lifespan and shows resistance to oxidative stress (Mourikis *et al*. 2006). These findings suggest that mitochondrial lipid metabolism might be important in proper mitochondrial function.

Both the *bss* and *jus* mutants showed some but not all of the characteristics associated with age-related dysfunction. *bss* showed no reduction in lifespan, although the population did show significant worsening of phenotype and a clear deficit in climbing ability. The *bss* mutant encodes an allele of the sodium channel and is probably not impacting mitochondrial function, although it does share some of the same characteristics as the *tko* and *ses B* mutants. The decline observed in the phenotype with age could be due to a general impact of aging on the nervous system. For example, wild type flies have to observe to develop temperature sensitive paralysis with age and bang-sensitivity is sometimes seen in our hands in very old wildtype flies (Reenan and Rogina 2008, personal observation). The impact of melatonin then could be a general impact on aging in *bss*. The *jus* mutant flies did not show many characteristics seen in other bs mutants. The lifespan was not significantly decreased and the recovery time remained unchanged throughout lifespan. Interestingly, *jus* did show a small decrease in climbing ability with age, although early testing showed climbing ability that was equivalent to the wild type. The results presented here suggest that the bang-sensitive phenotype must arise from a different pathway in these mutant. Further research must be done to find the link between these and the mutants that impact mitochondrial function

ROS damage in these mutants could be responsible for some of the age-related phenotypes. Our data suggests the all mutants with lifespan and age-related phenotypic decline except *ses B* respond to melatonin, a known free-radical scavenger. Since *ses B* has the greatest impact on energetics, its defects may not be amendable to rescue. Previous studies have shown the long-term supplementation of melatonin increased the lifespan of wild type *Drosophila* by decreasing the amount of unreduced oxygen species within the flies (Bonilla *et al*. 2002. We see a slight lifespan extension with wildtype in this study, although the effects on lifespan are more profound in *tko* and *eas*). Sanz *et al*. (2010) have showed that mitochondrial ROS production correlates with age, but its role in lifespan has been debated.

Alterations in neural function and reduced climbing ability with age have been correlated with neurodegeneration in the *Drosophila* (LeBourg & Lints 1991; Fergestad *et al*. 2006a; Reenan and Rogina 2008). However, there has been some variability in these findings. For example, flies with mutations in K+ channel genes are characterized by increased excitability, neurodegeneration and shorter lifespan in some studies (Trout & Kaplan 1970; Rogina & Helfand 1995) but not others (Fergestad *et al*. 2006b). Fergestad *et al*. (2008) reported that bs mutants have a reduced lifespan and exhibit neurodegeneration. Their lifespan findings are somewhat different from this work even with a similar dietary formulation: severe lifespan defects are observed with *bss* rather than the mitochondrial mutants *tko* and *ses B*, and they did not find a lifespan defect for *eas*. Fergestad *et al*. (2008) also observed age-dependent neurodegeneration that they described as correlated to in severity to seizure susceptibility and phenotype. In addition, using double mutants between *eas* and *ses B*, they found that there was a strong correlation between the seizure and neurodegeneration phenotypes. Our data suggests that bs mutants may have different paths to bang-sensitivity and age-related phenotypes, and in fact the phenotypes are separable. In our studies, age-related phenotypes are not consistently seen with all the bs mutants and are not correlated with bs phenotype.

In mammalian systems, epilepsy has been associated with neurodegeration in some cases, but not others. Temporal lobe epilepsy, the most common form, is associated with degeneration of the hippocampus. However, in a rat model, neuronal loss is intensified or increases severity of seizures, but was not responsible for their genesis (Zhang *et al*. 2002b). Kainate acid induced rats showing seizure do not have neuronal loss (Sloviter 1992; Buckmaster and Dudek 1997). AED have been associated with neurodegeneration at human treatment levels in young rats and AEDs have not been shown to rescue neurodegeneration associated with epilepsy (Bittigau *et al*. 2002).

Our data points to areas where the bs mutants have similarities and differences. We find that mutants associated with mitochondrial function have the most severe age-related phenotypes as expected, even if the strength of their excitability phenotypes are variable. To some extent, antioxidants can rescue age-related phenotypes, except in the case of *ses B* where viability may be challenged by low ATP levels. Treatments that increase or decrease excitability do not impact these age-related phenotypes, suggesting that they may arise through the mutant’s impact on pathways that are independent of the excitability defect.

## ACKNOWLEDGEMENTS

This work was completed with many Lafayette undergraduates contributing small parts to the larger work. The undergraduates who contributed are: Brooke Keim ’03, Kyle Klitsch ‘04, Erin Wolfson ’05, Lisa Consensa ’05, Diana Crai ’06, Blaine Caslin ’13, Eugene Warnick ’15, Stacy Nganga ’16, Dan Dellovade ’16, Ashley St. John ’18. Stephanie Cotes also contributed as a technician (NSF 2006).

